# Carrot storage diseases predominantly caused by mixed fungal infections

**DOI:** 10.64898/2025.12.09.693187

**Authors:** Karisto Petteri, Latvala Satu, Haapalainen Minna, Velmala Sannakajsa, Pennanen Taina, Suojala-Ahlfors Terhi

**Affiliations:** Natural Resources Institute Finland, Tietotie 4, Jokioinen, Finland; Natural Resources Institute Finland, Latokartanonkaari 9, Helsinki, Finland; University of Helsinki, Latokartanonkaari 7, Helsinki, Finland; Natural Resources Institute Finland, Toivonlinnantie 518, Piikkiö, Finland

## Abstract

Fungal pathogens often cause severe post-harvest losses in stored carrot yields. In 2020 and 2021, carrot samples were collected in Finland from 26 fields with different soil types. The proportion of symptomatic carrots was assessed at two time points during the cold storage period. To analyse the occurrence of different fungal pathogens in the symptomatic carrots, DNA samples were prepared and tested for the presence of fungal pathogens by species-specific PCR. The most frequently detected pathogens were *Cylindrocarpon* spp. with 61% occurrence, followed by *Fusarium avenaceum* (44%), *Mycocentrospora acerina* (42%), *Botrytis cinerea* (12%), and *Alternaria* spp. (0.8%). *B. cinerea* and *F. avenaceum* were more common in carrots grown on organic soil type, while *M. acerina* was more common in carrots grown on mineral soils. Mixed fungal infections were overrepresented in the symptomatic samples, in particular, *Cylindrocarpon* spp. often occurred together with *F. avenaceum* and *M. acerina* in the same samples. Amplicon sequencing of the fungal internal transcribed spacer region from the DNA of symptomatic tissues revealed that in addition to the pathogens - *B. cinerea, M. acerina, Cylindrocarpon* spp. and *Sclerotinia sclerotiorum* - the hyphomycete fungi *Tetracladium* spp. and the yeast *Leucosporidium intermedium* were abundant, and that *Tetracladium* spp. were more abundant in the carrots infected with *M. acerina*. A pathogenicity test confirmed that *F. avenaceum* is pathogenic to both the new season and old season carrots.

## Introduction

Pathogens spoiling carrot during long cold storage were recently surveyed in Finland, and *Mycocentrospora acerina, Botrytis cinerea, Fusarium* spp. and *Cylindrocarpon* spp. were identified as the major fungal pathogens (Latvala et al., 2024). In comparison to the earlier studies carried out in Finland (Parikka, 2008; Suojala, 1999), an increasing role of *Fusarium* spp. as pathogens of carrot was noticed. While the *Fusarium* spp. could be detected due to their high prevalence and compatibility with the culturing method used, some minor and/or slow-growing pathogens might still have remained undetected by this method. Possibly, minor fungal pathogens could be detected by DNA-based detection methods, like PCR and amplicon sequencing, that allow simultaneous detection of several species from the same sample.

As both abiotic and biotic factors are variable in the environment, the quantitative balance between the microbial species and the role of each individual species within the community in plant roots may change in time (Huang et al., 2020). For plant health, one major cause of concern is whether the endophytic microbes or so-called secondary pathogens could aggravate the disease symptoms. Changes in the abiotic environment, new host plant species introduced, or the old host plants weakened by stress may even result in the usually non-pathogenic fungi turning pathogenic (Stanosz et al., 2001).

It has been a common practice in plant pathology to study each pathogen separately, in attempt to associate each disease with a single causal pathogen. However, recent studies suggest that similarly to other communities and ecosystems formed by multiple species, the interactions between different microbial species are of major importance also in the pathosystems. The interactions between plants and several different microbes may promote plant health, for example in case of antagonism (Patkowska et al., 2017), or lead to a more severe disease (reviewed by Lamichhane and Venturi, 2015). Mixed infections in plants have been mostly studied in relation to mixes of different strains of the same pathogen species and hence higher potential for genetic recombination. Simultaneous infections with several different virus species have also been studied, and in some cases, the symptoms were found to be more severe than the infections with a single virus species (Mukasa et al., 2003; Yoshida, 2020). Cases of simultaneous infection by two different bacterial pathogens in carrot have also been detected (Munyaneza et al., 2011). Recently, more attention has been paid to mixed infections and their consequences also in the case of fungal diseases of plants (Bernasconi et al., 2023), although examples of those cases have been reported earlier (Linde et al., 2002).

Traditional approaches, like isolating pathogens by culturing, have mostly aimed to isolation of the dominant pathogen. While the culturing methods are useful for isolating the major pathogen or the pathogen of interest from a symptomatic sample by using selective media, these methods may fail to reveal mixed infections, for example, due to a substantial difference in either the relative amounts of the different pathogens in the sample or in the growth rates of the pathogens *in vitro* on the chosen media. This masking effect may lead to underestimation of pathogen prevalence. Recent advances in molecular methods, like species-specific PCR and microbiome analysis, based on amplicon sequencing, have enabled detection of mixed infections directly from the plant specimens.

The present study was conducted to investigate the presence of disease symptoms and fungal pathogens in carrot yield from commercial fields. To achieve this, a combination of visual inspection, and PCR and sequencing methods was employed. The primary aims of this study were to determine the relative abundance of known fungal pathogens of carrot – including cases of mixed infections, and to identify potential novel and/or minor pathogens that may have been overlooked in the previous surveys. In addition, the interrelations between different fungal pathogens were studied.

## Methods

### Carrot material and field sampling

Carrot samples were collected manually in September–October, as close to the farmer’s harvest time as possible. In 2020, 14 fields were sampled between 15 September and 7 October, and in 2021, 12 fields were sampled between 22 September and 5 October. Three fields in 2020 and three fields in 2021 were under organic management. Carrots were collected from six sampling sites in each field, with 30-100 meters distance between the sites. In total 26 fields were sampled at 156 sampling sites. Approximately 16 kg of carrots were collected at each sampling site and subsequently divided to two 8 kg subsamples that were packed in polypropylene bags. All the sampled carrots were stored at 0−1°C. Details of the collection and field sites are presented in more detail by Velmala et al. (2025) who analysed soil and rhizosphere microbiomes and their connection with carrot storage losses.

### Evaluation of disease symptoms

The stored samples were inspected for disease symptoms at two time points, with a 6-week interval, in January and March, and the sample weight was measured before and after the storage period (storage losses were analysed by Velmala et al. (2025) focusing on the latter March samples). The symptomatic carrots were washed and divided into four categories based on the visible (macroscopical) symptoms, as described in Latvala et al. (2024):

1. Decay at the root tip.
2. Pits and cavities on the side of the root.
3. Black or dark brown rot in the crown of the root.
4. Other symptoms, such as carrots spoilt by grey mould or Sclerotinia rot, with the pathogen identifiable by sclerotia, or other type of rot symptoms.

### Analysis of the symptomatic carrots

#### DNA extraction and species-specific PCR

In 2020, samples of symptomatic carrot tissue were taken in January and March from 181 individual lesions or spots on carrots from five different fields for identifying the pathogens. Symptomatic carrots were selected for PCR testing and sequencing from plots that differed in terms of carrot storage stability, plot soil type, cultivation method, and carrot cultivation history. Four samples belonging to the same symptom category were tested from each of the six subplots of the plot, i.e. up to 24 samples per symptom category per plot, if that many samples were available. Carrots of each symptom type in a plot/subplot were selected to represent the typical symptoms as well as possible. Of the tissue samples, 53.5% were of the root tip symptoms, 41.7% of pits on the side of the root and 4.8% of the crown black rot. The tissue samples were stored frozen at −80 °C prior to DNA extraction. Additionally, eight visually healthy individual carrots were sampled. Then 0.1 g of each tissue sample was ground using FastPrep-24 tissue and cell homogenizer (MP Biomedicals) in tubes containing Lysing matrix A (MP Biomedicals). The total DNA was extracted from individual samples using DNeasy Plant Mini Kit (Qiagen).

The DNA samples were tested for the presence of *M. acerina, F. avenaceum, B. cinerea, Alternaria* spp. and *Cylindrocarpon* spp. complex by single PCR reactions using genus or species-specific primers. The samples positive for *Alternaria* spp. were subsequently studied by sequencing the PCR product obtained with primers specific for the *Alternaria* major allergen *alt a1* gene region (Pavón et al., 2010). The PCR primers, annealing temperatures and the sizes of expected PCR products are shown in Table 1.

**Table 1.**
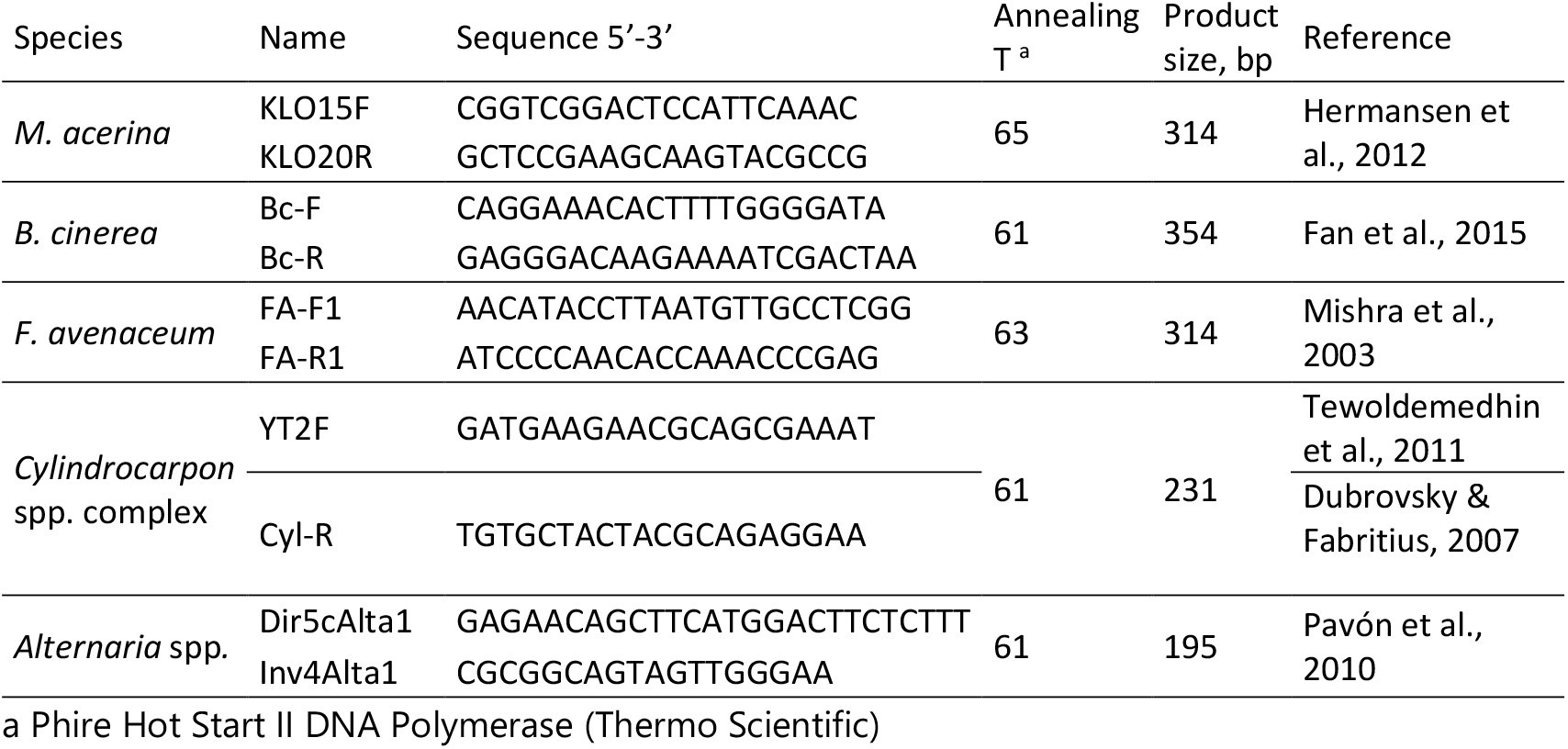
PCR primers used for detecting fungal pathogens at species or genus level.

PCR was run using 0.4 μL Phire Hot Start II DNA polymerase with its 1x reaction buffer (Thermo Scientific), 200 μM dNTPs, primers at 250 nM concentration, and 20 ng template DNA in 20 μL reaction volume. The programme was as follows: 98°C for 30 s; 35 cycles of 98°C for 10 s, annealing for 10 s (temperatures in Table 1), extension at 72°C for 30 s; followed by a final extension at 72°C for 5 min. The PCR products were analysed by electrophoresis on 1.5% agarose gels in Tris-borate-EDTA buffer. Gels were stained with ethidium bromide and visualized under a UV-transilluminator (SCIE-PLAS).

In 2021, tissue samples were taken from different types of symptoms in carrots originating from 9 different fields. In addition to 279 symptomatic carrots, DNA extraction and molecular analyses were performed with 13 healthy carrots. In January, the tissue samples were first cut in half. One half was used for DNA extraction on its own, whereas the other half was combined with other samples taken from similar symptoms of carrots from the same field plot and this mixture was used for DNA extraction. Altogether, 40 pools of symptomatic samples and 4 pools of healthy samples were constructed, each pool containing 2-6 individual samples. In March, a separate DNA sample was prepared of every sample, to be tested by PCR.

For the symptomatic samples from January, PCR tests were first performed on the pooled samples, to detect the five fungal pathogens named above. If a pooled sample was positive for any of those pathogens, then the corresponding single samples were tested separately with the same PCR test. The PCR reactions were performed as described above. For the individual samples from March 2021, only four pathogens were tested, excluding *Alternaria spp*., which had been rarely detected in January 2020 and not detected at all in the January 2021 samples.

#### Sequencing & bioinformatics

The 40 pooled DNA samples from symptomatic carrots and four pooled samples from healthy carrots from 2021 were further analysed by amplicon sequencing. Additionally, 11 individual symptomatic samples and three healthy samples from January 2020 were sequenced. Amplification of DNA from the pooled samples and amplicon sequencing, using Illumina MiSeq V3, paired end 2 × 301 bp, 2×10bp dual index, MCS 2.5.0.5/2.6.2.1 and RTA 1.18.54.0, were done at the Institute of Genomics, University of Tartu, Estonia (Caporaso et al., 2012). For amplification of the fungal internal transcribed spacer region ITS2 primers gITS7 (Ihrmark et al., 2012) and ITS4 (White et al., 1990) were used. The bioinformatic analyses were conducted with a multi-step pipeline as described in detail by Velmala et al. (2025; the parts focusing on the fungal sequences).

### Pathogenicity test with *Fusarium avenaceum*

Adapting a previously published method (Stanković et al., 2015), pathogenicity of *Fusarium avenaceum* FAV11, previously isolated from a symptomatic carrot in Finland (Latvala et al., 2024), was tested on both fresh new season carrots and old season carrots stored over winter. Carrots were first washed with water, then surface-sterilized in 0.05% sodium hypochlorite solution for 10 min, rinsed twice with reverse osmosis water and gently wiped dry. From each of the washed carrots, a small piece of peel was removed with a sterilized scalpel. For both new and old season carrots, 10 roots were inoculated by placing a 4 mm wide piece of the *F. avenaceum* FAV11 culture, grown on potato dextrose agar (PDA), upside-down at the wound site. In the control treatment, a sterile piece of PDA was placed on the wound site. Both the inoculated and control carrots were placed on sterile cloth inside plastic boxes, with the inoculation site upwards, and incubated in darkness at room temperature (22 °C). The width and height of the visible symptom and/or fungal growth were measured with a digital caliper, 7 and 18 days after the inoculation.

### Statistical analysis

Data analysis was implemented mainly in Python 3.12. Package *pandas* v2.2.3 (McKinney, 2010) was used for data curation. Figures were created using packages *matplotlib* v3.10.1 (Hunter, 2007) and *seaborn* v0.13.2 (Waskom, 2021). Symptom and pathogen distributions and relationships were tested with Kruskal-Wallis test as function Kruskal, chi-squared test using function chi2_contingency, and Pearson correlation with function pearsonr in package *scipy* v1.15.2 (Virtanen et al., 2020). Pairwise associations between species occurrence in the PCR data were additionally tested in R (R Core Team, 2024) using package cooccur (Griffith et al., 2016). Pairwise correlation between ITS-read counts among top 15 taxa were analysed with Pearson correlation (pearsonr) using adjusted p-value based on Bonferroni correction for multiple testing.

The python source code for data-analysis and the data is deposited in a Zenodo repository 10.5281/zenodo.17212592 (will be made public upon publication). Raw amplicon sequence data are deposited in the sequence read archive SRA of the NCBI database under BioProject PRJNA976070 with accessions SAMN35358255-SAMN35358630.

## Results

### Carrot health

Altogether, 2,500 kg of carrots were taken into storage, and all the 32,815 individual carrots were visually inspected for disease symptoms after the storage period (Table 2). Of all the inspected carrots, 16.6% (5447 pcs.) were symptomatic. The most common symptom class was the root tip symptoms (65.4%), followed by side symptoms (28.2%) and crown symptoms (3.1%). Other symptom types, such as (an extensive symptom of) Sclerotinia rot or grey mould, appeared in 3.4% of the symptomatic carrots.

**Table 2.** Summary of carrot sampling and analysis.

| Time | # Fields (# organically managed) | Total carrots | Symptomatic carrots | Fields for DNA samples | DNA samples (symptomatic) | DNA samples (healthy) | PCR Tests | NGS |
| --- | --- | --- | --- | --- | --- | --- | --- | --- |
| 2020 J | 14 (3) | 8144 | 1123 (14%) | 5 | 77 | 8 | 5 pathogens | 11 symptomatic + 3 healthy samples |
| 2020 M | 14 (3) | 8276 | 2073 (25%) | 3* | 104 | 0 | 5 pathogens | - |
| 2021 J | 12 (3) | 8065 | 729 (9%) | 9 | 176 (into 40 pools) | 13 (into 4 pools) | 5 pathogens (pools first) | 40 symptomatic + 4 healthy pools |
| 2021 M | 12 (3) | 8330 | 1522 (18%) | 3* | 103 | 0 | 4 pathogens (excl. *Alternaria*) | - |
| **Total** | **26 (6)** | **32815** | **5447 (17%)** | **14** | **460** | **21** |  |  |

\* The samples in March (M) 2020 and March 2021 were taken from the same field batches sampled in January (J) of the same year.

Storability of the carrots varied between the fields and the years (Figure 1). At the latter time point, in March, the mean proportion of symptomatic carrots was significantly higher in 2020 (26%) than in 2021 (18%, p ≈ 0.0003, Kruskal-Wallis). Carrots from the fields with organic soil type had a mean symptom rate of 26% over the two years, which was significantly higher than the 18% symptom rate of carrots from the fields with mineral soil type (p ≈ 0.0004). Within the individual years, the difference between organic and mineral soil type fields was significant only in 2021 (p ≈ 0.0005).

**Figure 1.**
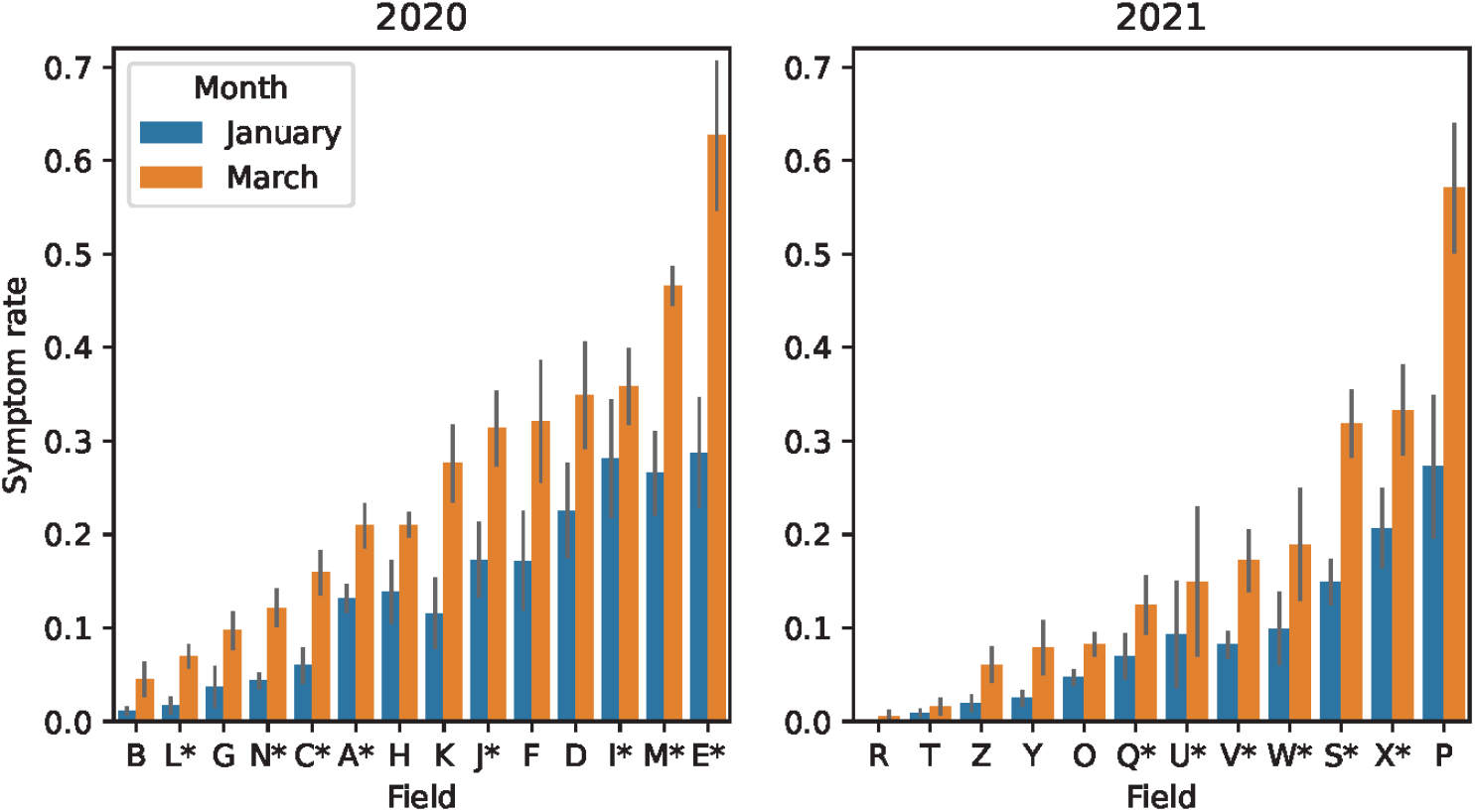
Proportion of symptomatic carrots in samples from different fields in January (blue) and March (orange). The mean symptom rates for six samples per field are shown. The error bars denote standard error of the mean. Fields with organic soil type (>20% organic matter) are marked with asterisk after the code letter.

Samples from each symptom class, altogether 460 individual symptomatic samples, and 21 healthy samples were selected for pathogen screening by specific PCR primers. The most frequently detected pathogen was *Cylindrocarpon* spp. with 61% occurrence, followed by *F. avenaceum* (44%), *M. acerina* (42%), *B. cinerea* (12%), and *Alternaria* spp. (0.8%). The samples that were positive for *Alternaria* spp. were analysed by sequencing the PCR product. Two of the samples yielded readable sequence: one was identical with *A. alternata*, and the other matched several different species including for example *A. atra* and *A. consortialis*, making the species level identification inconclusive. The sequences were deposited to GenBank (accession numbers PV448265 and PV448266). Based on the sequence, the species *A. radicina* could be excluded. On average, 1.60 different pathogens were detected per one symptomatic sample. Of the symptomatic samples, 128 (28%) were negative for all tested fungal pathogens.

### Pathogens across symptom types

The presence of screened pathogens was analysed in different symptom types, to detect any differences in the preferred site of infection between the different pathogens. The majority of the tests were performed on root tip symptoms (246, 53%), followed by side symptoms (192, 42%), and crown symptoms (22, 5%). No major differences were observed in the pathogen distribution between the different symptom types; instead, the frequencies reflected well the pathogens’ overall occurrence, although the test indicated nearly significant differences (p-value: 0.056, chi-squared). However, *Cylindrocarpon* spp. seem to occur in the side symptoms less frequently than expected. *B. cinerea* was not detected in any of the crown symptoms, while *Alternaria* spp. were not detected in the side symptoms. Especially with *Alternaria* spp. the low overall frequency is likely to explain their absence from one of the symptom classes. Similar argument might explain the absence of *B. cinerea* in the least common symptom type (only 22 crown samples were tested). In general, the pathogen distributions were quite similar between the three classes of symptoms (crown, side and tip) (Figure 2).

**Figure 2.**
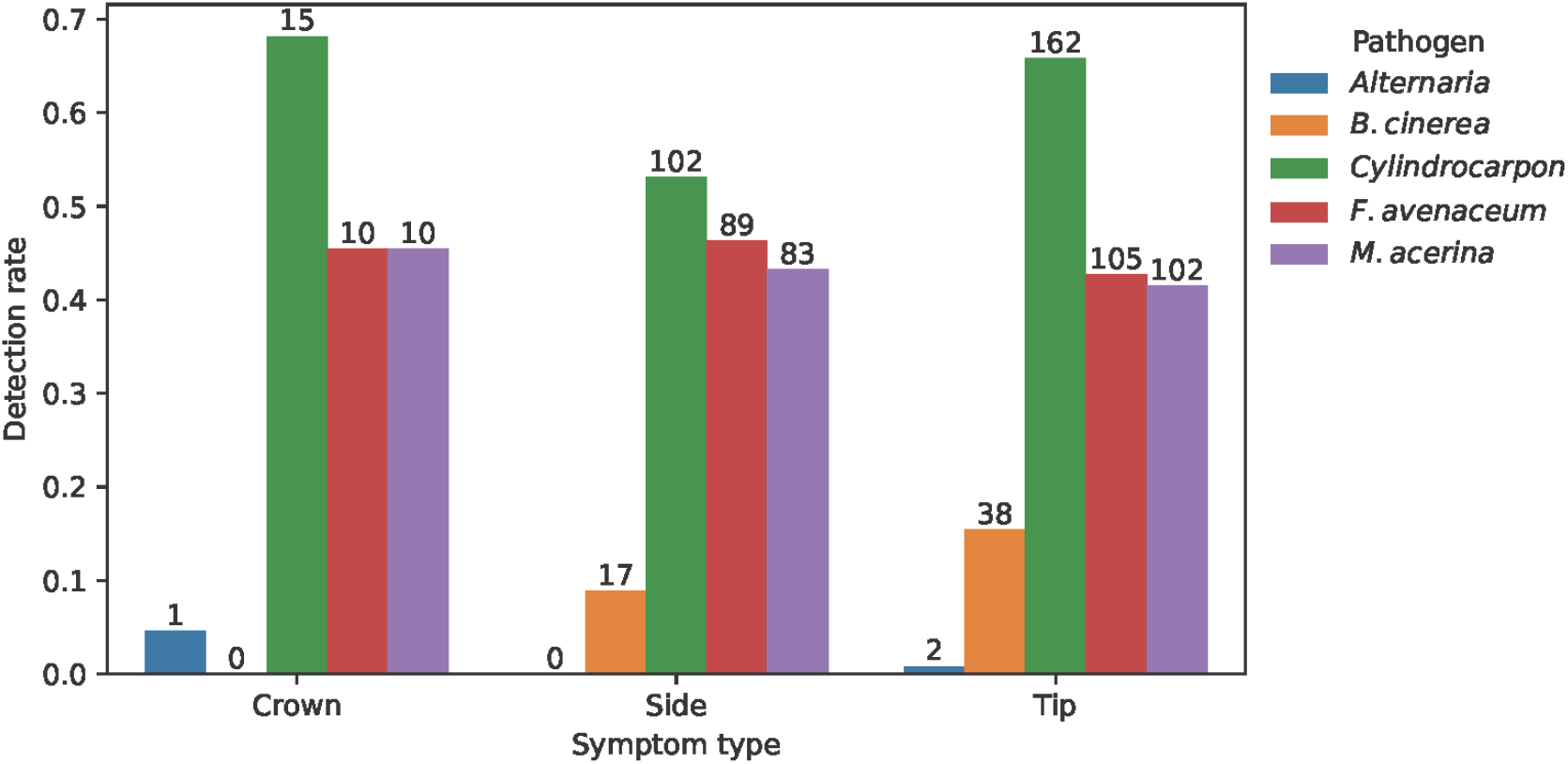
Prevalence of the five pathogens in different symptom types based on PCR testing.

### Pathogens in different soils

The symptomatic samples identified with PCR represented equally the growing sites with either organic or mineral soil type (230 samples each). Significant differences were observed in the distribution of different pathogens between the two soil types (p-value: 5.3*10^-5, chi-squared). Especially, *B. cinerea* and *F. avenaceum* were more common in the organic soil type, while *M. acerina* was slightly more common in mineral soil type (Figure 3).

**Figure 3.**
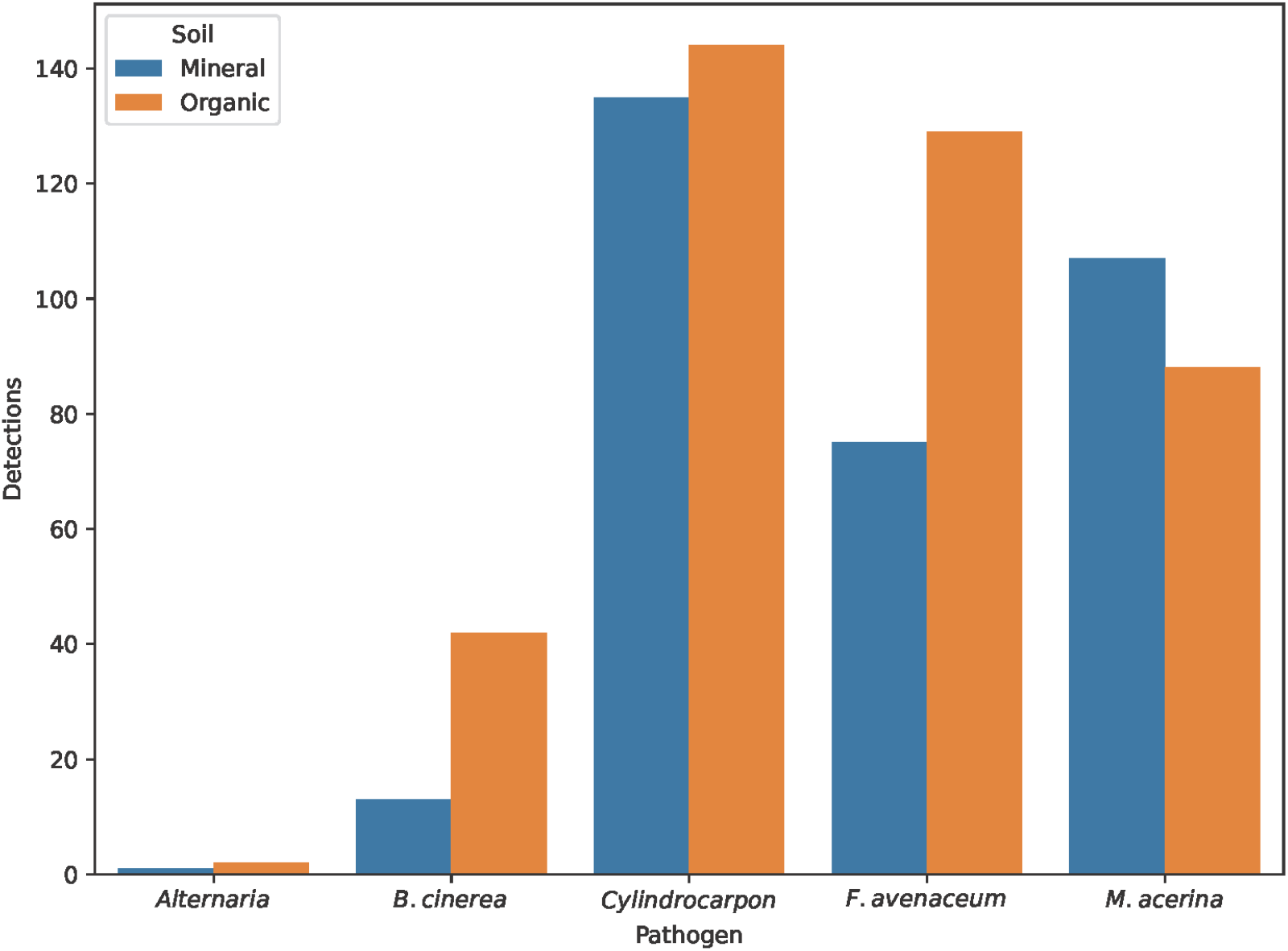
Observed pathogen distributions in the symptomatic samples from carrots originating from fields with mineral (blue) or organic soil type (orange). B. cinerea and F. avenaceum were more common in carrots from organic soil fields, whereas M. acerina was more common in carrots from mineral soil.

### Co-occurrence of pathogens in individual symptoms

As implied by the high average pathogen count per symptomatic sample (1.60), most of the samples contained multiple pathogens. Comparison of the expected counts of pathogen combinations to the observed counts revealed that mixed infections were indeed overrepresented. First, samples with no detected pathogens were overrepresented more than two-fold compared to expectation, indicating that pathogens appeared together more often than expected (Figure 4). Second, all single infections were observed less frequently than expected, especially in the case of *Cylindrocarpon* spp. In turn, *M. acerina, F. avenaceum, B. cinerea* and *Cylindrocarpon* spp. seem to prefer each other, since the two triplets MFC and FBC, and all four together (MFBC) were clearly overrepresented. Interestingly, *B. cinerea* was not detected alone nor in combination with *M. acerina*, although these cases were expected to occur six and five times, respectively.

**Figure 4.**
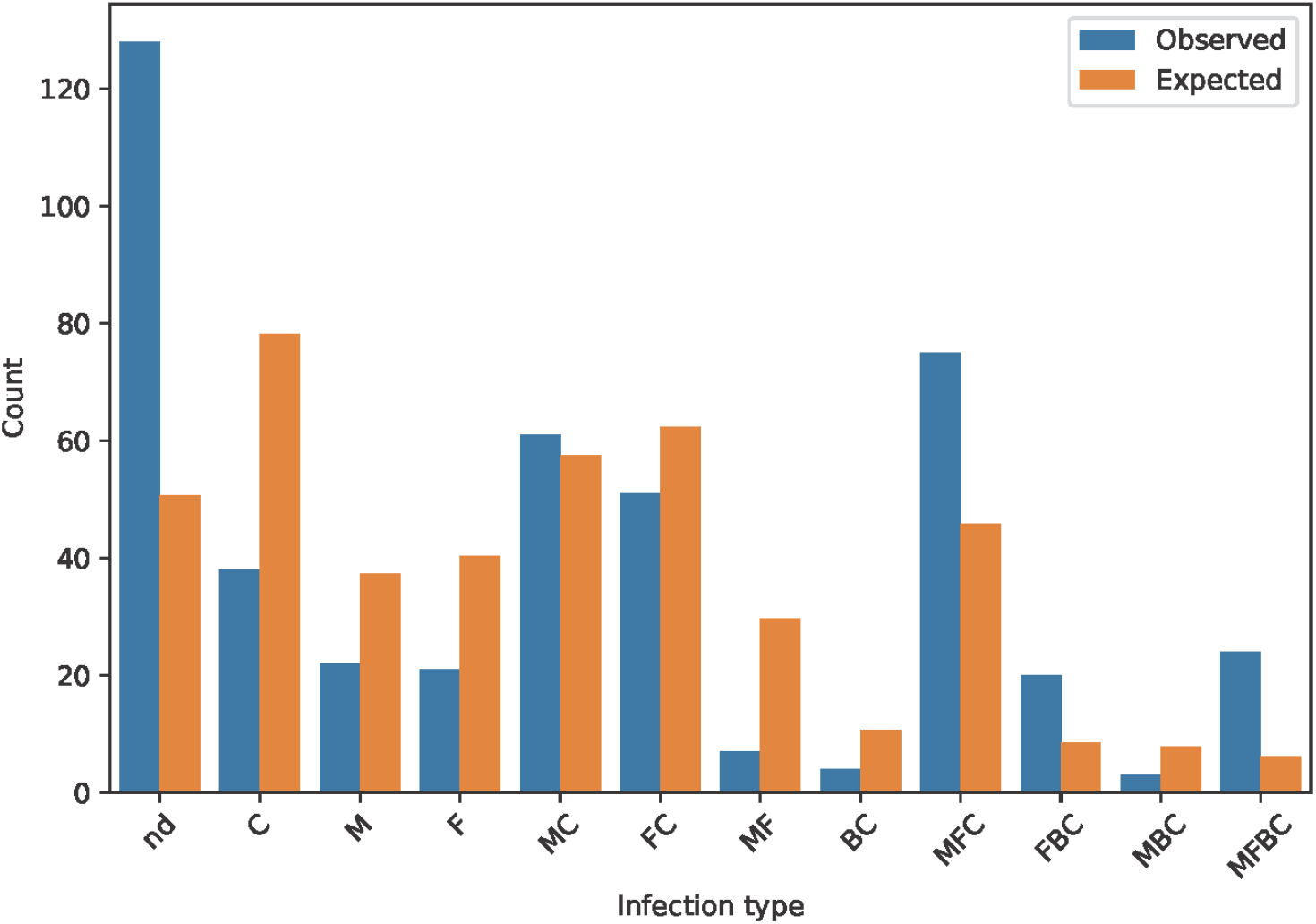
Number of different types of single-pathogen and mixed infections based on PCR testing (blue) compared with expected (orange) distribution of different types based on random occurrence of symptoms (letters refer to the first letter of the species’ name, nd=not detected). The rarest observed pathogen combinations are not shown in the figure. They were ‘FB’ (observed: 2, expected: 5.5), ‘MFB’ (o: 1, e: 4.0), ‘FAC’ (o: 1, e: 0.5), ‘MFAC’(o: 1, e: 0.4), ‘MFBAC’(o: 1, e: 0.1). The non-observed combinations are likewise not shown. They were expected less than once except for B (e: 6.9) and MB (e: 5.1).

The pairwise co-occurrence analyses revealed in total five significant associations (co-occurrences). These were observed between *Cylindrocarpon* spp. and either *M. acerina* (p-value <10^-5, cooccur), *F. avenaceum* (p-value <10^-5), or *B. cinerea* (p-value <10^-5), and additionally between *F. avenaceum* and either *M. acerina* (p-value: 0.0003) or *B. cinerea* (p-value <10^-5). The strongest associations were observed between *Cylindrocarpon* spp. And *M. acerina* (observed together: 165, expected together: 118), and between *Cylindrocarpon* spp. and *F. avenaceum* (observed: 173, expected: 124)

### Sequencing results

Sequencing the 11 individual samples and 40 pools of symptomatic carrot tissue together with three individual healthy samples and four healthy pools confirmed the previously identified pathogens and revealed other fungal species not associated with carrot diseases. In addition to the known carrot pathogens - *Botrytis cinerea, Mycocentrospora acerina, Cylindrocarpon* spp. and *Sclerotinia sclerotiorum* - sequencing revealed the presence of hyphomycete fungi *Tetracladium* spp. and the yeast *Leucosporidium intermedium*.

The most common taxa based on the relative read counts across all the samples were *B. cinerea* (33% of the reads), *M. acerina* (26%), *Tetracladium* spp. (20%), *Cylindrocarpon* spp. (4.8%) and *Sclerotinia sclerotiorum* (3,7%). *F. avenaceum* was only detected at 0.44% of the reads, contrasting with its high prevalence according to PCR, and *Alternaria spp*. were not among 50 most common taxa (less than 0,1% of the reads). Some additional interesting taxa were detected: unidentified fungi belonging to the order *Helotiales* (potentially *Cadophora* spp.) were the sixth most common (3,4%), and *Plectosphaerella cucumerina* was the 11^th^ most common (0.47%).

Correlation analysis between the number of sequencing reads and the positive PCR results across the individual samples (opened pools) revealed that the results obtained with the two methods were not strongly connected. Among the four major pathogens detected by PCR, only *B. cinerea* showed significant Pearson correlation (r = 0.596; p=0.019) and *M. acerina* nearly significant (r=0.462; p=0.083). For the others the connections were non-significant (*Cylindrocarpon* spp. p=0.429; *F. avenaceum* p=0.488).

Five significant pairwise correlations were detected among the 15 most common taxa. These were observed between *M. acerina* and *Tetracladium* spp. (p-value: 0.000189); between *S. sclerotiorum* and *L. intermedium* (p-value: 1.0*10^-17); and between *Cladosporium* spp. and either *B. cinerea* (p-value: 0.000141) or *Helotiales* spp. (potential *Cadophora* spp.) (p-value: 0.000002) or *Pseudogymnoascus roseus* (p-value: 0.000017).

### Pathogenicity assay

In the inoculated carrots, *F. avenaceum* caused wide symptoms with white fungal mycelia on the top, and at the later time point the symptom extended around the whole root in most of the carrots. No symptoms were detected at the mock inoculation site of the control carrots. The carrots similarly inoculated with *Plectosphaerella* spp. showed no symptom development (data not shown).

The old, overwintered carrots were more susceptible to *F. avenaceum* infection than the new season carrots, and a significant difference in symptom development was observed (two-sample t-test, p<0.01) between the carrot batches (Figure 5). In between the two time points, *F. avenaceum* was able to cause a progressive disease on both the old and the new season carrots (paired t-test, p<0.01).

**Figure 5.**
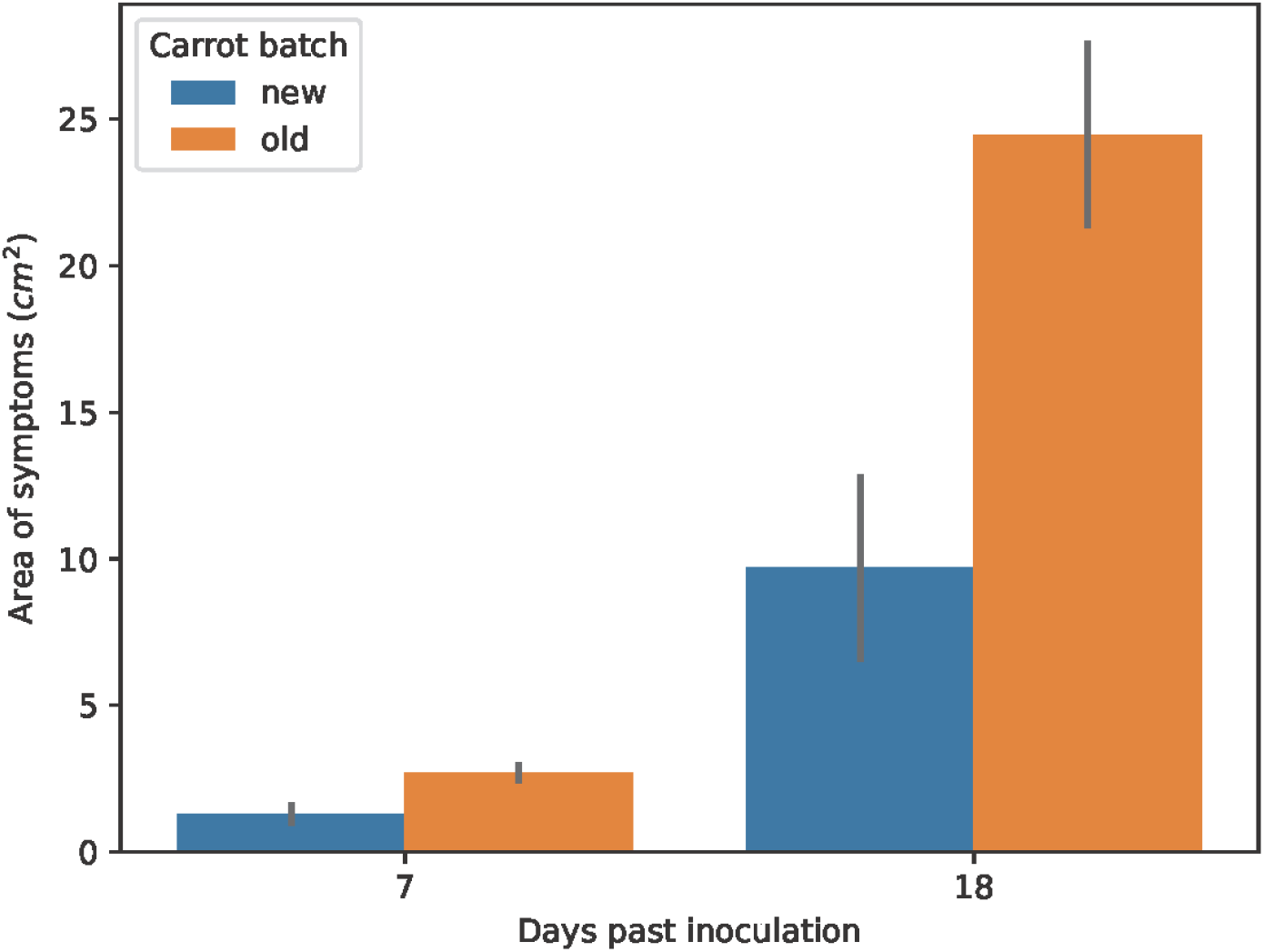
Area of symptoms on F. avenaceum inoculated carrots at two time points on two different batches of carrot. Error bars show SE.

## Discussion

We conducted a survey of diseases and fungal pathogens of carrot over several fields and two growing seasons, using visual assessment for disease evaluation, and taxon-specific PCR and ITS-sequencing for pathogen detection and identification. The pathogen prevalence was analysed in relation to the symptom type, the field soil type, and the presence of other pathogens. Interestingly, in the majority of those symptomatic samples in which pathogens were detected there were several different pathogen species. Single infections with only one pathogen detected were less common than expected based on random distribution. Hence, mixed infections seem to be dominant in carrot storage diseases.

The observed carrot storability varied significantly between field lots and years. Climatic and other abiotic factors affect carrot health (Goodliffe and Heale, 1977; Hermansen et al., 2000; Suojala, 1999) as do biotic factors such as field microbiome (Velmala et al., 2025). Intensive carrot cultivation with poor crop rotation increases post-harvest losses due to diseases (Latvala et al., 2024). The devastating scale of post-harvest losses, up to 60% in the worst cases, highlight the importance of understanding the factors affecting the occurrence of carrot post-harvest diseases. In addition to the cultivation practices and the weather during the growing season, also the soil type affects the occurrence of post-harvest diseases. The carrots grown in organic soils seem to have slightly higher storage losses due to diseases than those originating from mineral soil fields.

The most common pathogen identified by species-specific PCR in this study was *Cylindrocarpon* spp. with 61% occurrence. This result is different from the result previously obtained from the 2016-2018 sampling by culturing method (Latvala et al., 2024), in which *Cylindrocarpon* spp. were only detected in 4.7% and 6.0% of the carrot root tip samples in January and March, respectively. In this study, *Cylindrocarpon* spp. was detected by PCR in the symptomatic samples mostly together with the other fungal pathogens, which suggests that the culturing method underestimated the prevalence of this pathogen due to its lower competitiveness on the common culture media. The next most common pathogens were *F. avenaceum* and *M. acerina* at similar abundance (44%, 42%, respectively) and *B. cinerea* at lower prevalence (12%). This is quite similar pattern although at lower level compared to Latvala et al. (2024) who detected *M. acerina* and *Fusarium* spp. By PCR at high levels (>80%, ca. 70%, respectively) and *B. cinerea* clearly less commonly (<20%). Similarly in their culturing results, *M. acerina* was the most common fungal pathogen, followed by *Fusarium* spp. and *B. cinerea* (in the root tip samples). In contrast, *Alternaria* spp. were only detected in three samples in this study, and thus they seem to be rare in the carrot roots in Finland. Our further sequencing of the *Alternaria*-positive PCR products confirmed that none of the samples contained *Alternaria radicina*, a seed-borne pathogen that is a major concern in carrot and other apiaceous crops in many other countries (Farrar et al., 2004). Rarity of *Alternaria* spp. is supported by the aforementioned culturing results (2016-2018), where the taxa was likewise not detected (Latvala et al., 2024).

The majority of the symptoms were observed at the root tips, while crown symptoms were not common, similar to earlier results in Finland (Latvala et al., 2024). This might indicate the importance of the breakage of the tip of the tap root as the initial infection site or as a site suffering from loss of water and vigour. Similarly, root and petiole breakage might explain side and crown infections, as previously suggested for *M. acerina* (Davies et al., 1981). All the five fungal pathogens studied were observed fairly equally in all the symptom types, suggesting site-independent infection mechanisms. In contrast, there was a significant difference in the pathogen distribution between the organic and mineral soils. Most notably, *F. avenaceum* and *B. cinerea* were much more common in the carrots from organic soils, whereas *M. acerina* was more common in the mineral soils. This agrees with the previous results obtained in Norway, where the stored carrots grown in sandy soils had more liquorice rot than the carrots grown in peat soil (Hermansen et al., 2000).

Co-occurrences of *Cylindrocarpon* spp. and/or *Fusarium avenaceum* together with the necrotrophic pathogens *M. acerina* and *B. cinerea* were observed more frequently than expected. This suggests that in carrot *Cylindrocarpon* and *Fusarium* could often act as secondary pathogens and saprotrophs, taking advantage of the tissue damage caused by the necrotrophs. As the pair *M. acerina* and *B. cinerea* was not detected without those other fungi; the two necrotrophs may not benefit from each other but rather compete with each other. A considerable proportion of the symptomatic samples were negative in the PCR tests for the screened pathogens *Cylindrocarpon* spp., *F. avenaceum, M. acerina, B. cinerea* and *Alternaria* spp., which were expected to be the major fungal pathogens of carrot in Finland based the previous studies (Latvala et al., 2024). This naturally opens questions about potential unidentified pathogens that could have been involved in the symptom formation. Since only three of those symptomatic samples that were negative in all the PCR tests were included in the fungal ITS sequencing, the possibility of some other fungal pathogens in the rest of the PCR-negative samples cannot be excluded. It is also possible that the main pathogen in those symptomatic samples was not a fungal but a bacterial species. Previously, *Pseudomonas* spp., *Pectobacterium* spp., *Dickeya chrysanthemi*, and *Erwinia rhapontici* were shown to cause decay of stored carrots in Finland (Kahala et al., 2012). Also, in the recent study by Latvala et al. (2024), a large proportion of the carrot spoilage was caused by bacteria after the exceptionally warm growing season in 2018. Despite the amount of samples with no detected pathogen, the mean number of pathogens per sample was 1.6 highlighting the dominant role mixed infections in the carrot post-harvest diseases.

Pathogenicity of *Fusarium avenaceum* covers a wide host-range, including cereals and leguminous plants (Safarieskandari et al., 2021). Recently, *F. avenaceum* has been reported also as a pathogen of carrot in several European countries – in Serbia (Stanković et al., 2015), France (Le Moullec-Rieu et al., 2020) and Finland (Latvala et al., 2024). Here we further confirmed experimentally that a *F. avenaceum* isolate originating from a symptomatic carrot grown in Finland was indeed infective and highly virulent on carrot. We also observed a significant difference in the disease development between the new season and old season carrots, supporting the view that the carrots’ susceptibility to diseases increases with a long storage period (Goodliffe and Heale, 1977; Latvala et al., 2024).

The ITS sequencing results were to some extent in line with the PCR results of this study and also earlier studies, as the known pathogens were among the most abundant species based on the relative read counts. The fungal species most frequently identified in the symptomatic carrot samples by sequencing were *B. cinerea* (4^th^ in PCR) and *M. acerina* (3^rd^ in PCR), while *Cylindrocarpon* spp. that were most frequently detected by PCR were the 4^th^ most common in the sequences. Another common pathogen of carrot, *S. sclerotiorum*, was the 5^th^ most common in the sequences. Hence, these pathogens were dominating in the samples of symptomatic carrot tissue.

In agreement with the PCR results, *Alternaria* spp. were rarely appearing in the sequences. Unlike for the other pathogens, the agreement between the PCR results and ITS sequence counts was poor for *F. avenaceum*, since it was very common in PCR (2^nd^, 44% of the samples positive) but not prominent in sequences (13^th^). The suitability of standard ITS-sequencing for detection and identification of *Fusarium sp*. is known to be low (O’Donnell et al., 2022), which potentially explains this discrepancy. Further, the low or non-significant correlations between PCR-positivity rate and sequencing results signify that these two detection methods are not equivalent but rather complementing each other.

The strengths of PCR are its sensitivity and specificity, as it provides results on exactly what is being looked for, and it is possible to optimise the workflow for each target species. PCR also provides a simple quantitative measure of the incidence or prevalence of a pathogen. The specificity of the test, however, can also be its weakness, since the results only show what was looked for. Instead, amplicon sequencing with general primers provides a chance to detect multiple species at once and also to detect unexpected taxa. However, in amplicon sequencing the false negative rate is likely to be much higher than in the species-specific PCR, because the general primers do not have a good match with all the fungal taxa. The suitability of the standard ITS-primers varies between taxa also due to the different copy number of the target sequence (Ihrmark et al., 2012). Moreover, the variation within the sequence is also different between taxa, leading to a variable power in species identification. Naturally, both of these molecular methods also detect DNA from non-viable sources, and hence isolation and inoculation experiments to fulfil Koch’s postulates are still required for confirming causality (Byrd and Segre, 2016).

*Tetracladium* spp., order *Helotiales*, were found to be more abundant in the carrot root samples heavily infected with *M. acerina* than in the *M. acerina*-free samples. While many *Tetracladium* spp. are common in soil and decaying plant matter, contributing to the recycling of nutrients, some species are beneficial plant root endophytes, enriched in the roots in comparison to bulk and rhizosphere soil (Lazar et al., 2022). The necrotrophic pathogens induce cell death in the infected plant tissues (Choquer et al., 2007; Louarn et al., 2012). Thus, *Tetracladium* could temporarily benefit from the attack by these necrotrophs and use the decaying carrot tissues as food source.

Among other notable species detected from the symptomatic samples, *Leucosporidium intermedium* (Nakase & M. Suzuki), syn. *Sporobolomyces intermedius, Bullera intermedia*, is a saprotrophic yeast that can be found in decaying organic material in the soil. This species was first isolated from dead leaves of rice (*Oryza sativa*) in Japan (Nakase and Suzuki, 1986). *Plectosphaerella cucumerina* and *Plectosphaerella plurivora* are common necrotrophic fungi found in agricultural soils. These and other *Plectosphaerella* spp. can cause root rot and foliar wilt symptoms in a wide range of cultivated vegetables (Carlucci et al., 2012). However, *Plectosphaerella* spp. can also use nematodes as food source and thus diminish nematode pressure on crop plants, and thus they have even been suggested as biological control agents against potato cyst nematodes (Atkins et al., 2003) and the wide host range nematode species *Nacobbus aberrans* (Sosa et al., 2018).

In conclusion, the presented data indicates that carrot storage diseases are predominantly caused by mixed fungal infections rather than single pathogens. *Cylindrocarpon* spp. frequently co-occur with necrotrophs such as *Mycocentrospora acerina* and *Botrytis cinerea. Fusarium avenaceum*, a broad host-range pathogen, was confirmed as highly virulent, and observed host resistance decreased with storage duration. Storability exhibited variability between the lots and years, influenced by soil type, climate, and composition of the pathogen communities, with organic soils demonstrating higher levels of loss. The presence of discrepancies between PCR and ITS sequencing, particularly in the case of F. *avenaceum*, underscores the complementary nature of these methods and the limitations of the ITS-barcode region for specific taxa. The high prevalence of mixed infections and substantial post-harvest losses highlight the necessity for integrated management strategies and broader diagnostics to address both fungal and bacterial contributors of carrot spoilage.

## Acknowledgements

This project was funded by the Ministry of Agriculture and Forestry of Finland (VN/4602/2020), and Maiju and Yrjö Rikala’s Horticultural Foundation. We thank researchers Pirjo Kivijärvi and Minna Pirhonen for their valuable input in designing and executing this experiment. We are grateful for the technical assistance of Senja Tuominen and Senja Räsänen in the laboratory and Johanna Rihtilä in performing the storage experiments and for analyses of storage losses. The authors declare that they have no competing financial interests or personal relationships that could have appeared to influence the work reported in this paper.

